# Carbapenem-resistance *oprD* mutations reshape *Pseudomonas aeruginosa* host-pathogen interactions during infection

**DOI:** 10.1101/2025.10.24.684370

**Authors:** Pablo Laborda, Claudia Antonella Colque, Ruggero La Rosa, Søren Molin, Helle Krogh Johansen

## Abstract

Antibiotic resistance is a major global health threat. While its role in reducing drug susceptibility is well established, the broader consequences of resistance mutations on bacterial physiology and behavior during infection remain poorly understood. Carbapenem-resistant *Pseudomonas aeruginosa* is considered among the highest- priority bacterial threats, with resistance commonly driven by loss-of-function mutations in the carbapenem uptake porin OprD. Here we show that such mutations can arise in clinical isolates even without prior carbapenem treatment, indicating that their effects during infection extend beyond antibiotic resistance. Consistent with this, we found that *oprD* mutants exhibit enhanced early attachment to and translocation across airway epithelial barriers in infection models, an effect maintained across strains with distinct clinical genomic backgrounds and infection dynamics. Our findings indicate that loss of OprD alters the bacterial outer membrane charge and reduces mucus entrapment, thereby facilitating epithelial barrier colonization. Overall, these results illustrate how antibiotic resistance mutations can directly shape infection dynamics, extending their impact well beyond antimicrobial susceptibility.

## Introduction

Antibiotic resistance is one of the most important medical challenges of the 21^st^ century, undermining the effectiveness of current treatments and driving higher morbidity and mortality associated with infections worldwide ^1^. Gram-negative pathogens are of particular concern due to their intrinsic resistance mechanisms, low outer membrane permeability to antimicrobial agents, and ability to acquire additional resistance traits ^2^. Among them, *Pseudomonas aeruginosa* stands out as a highly adaptable opportunistic pathogen capable of causing a broad spectrum of infections, especially in immunocompromised patients and individuals with chronic lung diseases such as cystic fibrosis (CF), Primary Ciliary Dyskinesia (PCD) and Chronic Obstructive Pulmonary Disease (COPD) ^3–5^.

In chronic infections, *P. aeruginosa* often acquires mutations that reduce the efficacy of antimicrobial agents ^6,7^, a phenomenon typically attributed to the selective pressure of antibiotic exposure. Resistance to carbapenems, a last resort antibiotics, is of particular concern because it severely restricts therapeutic options ^8,9^, placing carbapenem- resistant *P. aeruginosa* among the highest-priority bacterial threats ^10^. A central determinant of carbapenem susceptibility is the outer membrane porin OprD, which mediates drug uptake of carbapenem antibiotics. Loss or alteration of OprD is among the most common mechanisms of carbapenem resistance in *P. aeruginosa* ^4,11,12^, and it is frequently observed in clinical isolates ^6,7^.

While the genetic basis of antibiotic resistance is generally well-characterized, its broader physiological consequences remain largely undiscovered. Resistance mutations can exert pleiotropic effects on bacterial envelope organization, metabolism, and virulence ^13–17^. These changes may occur directly, when the altered resistance determinant is involved in core physiological processes, or indirectly through perturbation of complex regulatory networks. Notably, many of these networks are intertwined with virulence and pathogenicity pathways ^18,19^, suggesting that antimicrobial resistance can reshape infection dynamics in ways that extend beyond drug susceptibility.

Here, we show that loss-of-function mutations in *oprD* appear in clinical isolates in the absence of prior carbapenem exposure, suggesting that they can provide infection- related advantages beyond antibiotic resistance. Consistent with this, we observed that these mutations increase *P. aeruginosa* attachment to and crossing of airway epithelial barriers. Our data suggest that loss of OprD leads to a reorganization of the bacterial surface and reduces mucus entrapment, thereby facilitating earlier access to the epithelial surface and more rapid penetration. Together, these findings highlight how examining antimicrobial resistance and bacterial physiology as interconnected traits can reveal important links with direct consequences for infection outcomes.

## Results

### Mutations in oprD appear in clinical isolates without prior carbapenem exposure but often disappear

We first investigated the clinical prevalence of *oprD* mutations in chronically infected patients. For this, we examined a collection of 474 longitudinally sampled clinical isolates of *P. aeruginosa* obtained from 34 chronically infected individuals with CF at the Copenhagen CF Center ^6^. Most mutations in *oprD* acquired during infection were frameshifts, likely preventing porin production or assembly into the membrane (Figure 1A, Supplementary Table 1). These mutations were observed at multiple timepoints throughout chronic infection. When comparing these data with carbapenem treatment periods, we found that the appearance of *oprD*-mutated strains did not always coincide with antibiotic exposure (Figure 1B). Importantly, carbapenem administration in these patients was exclusively intravenous, as recorded in hospital treatment records, ensuring that treatment histories are not confounded by undocumented oral use at different timepoints. Consistent with this, meropenem Minimal Inhibitory Concentration (MIC) testing showed that a substantial fraction of *oprD*-mutated isolates (26%) remained susceptible according to EUCAST clinical breakpoints ^20^ (MIC ≤ 2; Supplementary Table 1), with susceptible isolates recovered both after carbapenem therapy and from patients without prior exposure. Together, these observations indicate that the emergence of *oprD* mutations is not solely driven by carbapenem pressure and may reflect infection-related advantages beyond antibiotic resistance.

**Figure 1.**
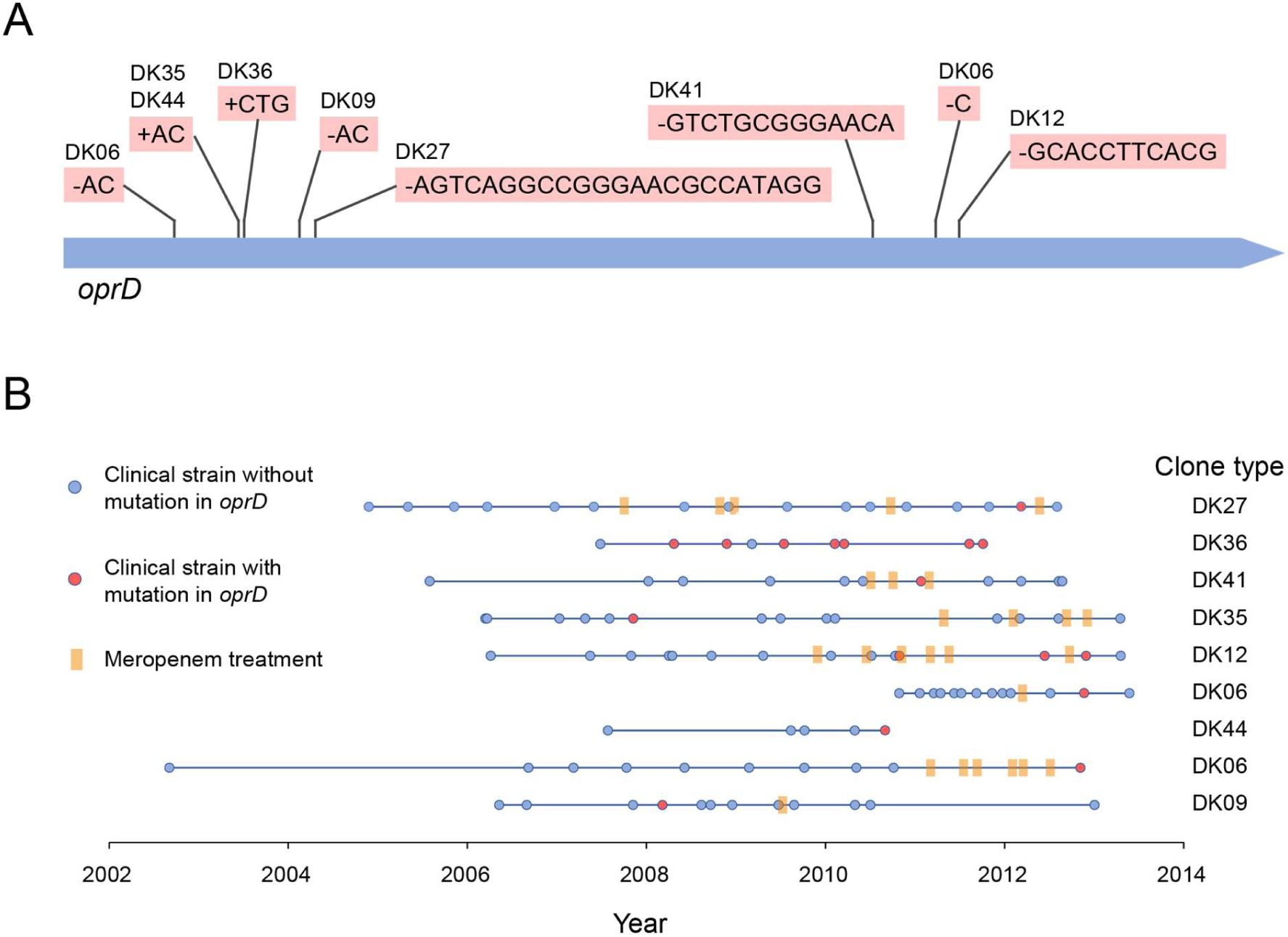
Characterization of *oprD* mutations in *P. aeruginosa* clinical isolates. (A) Non-synonymous mutations identified in the *oprD* gene of *P. aeruginosa* clinical isolates from chronically infected individuals with CF, belonging to different clone types ^6^. Information about the specific position of each mutation can be found in Supplementary Table 1. (B) Temporal distribution of *oprD*-mutated strains during chronic infection, shown together with carbapenem treatment periods of the corresponding patients. Isolates without mutations are indicated by blue circles, isolates carrying *oprD* mutations by red circles, and carbapenem treatment periods by orange rectangles.

Despite their recurrent acquisition, *oprD* mutations were almost never maintained in the infecting populations of this cohort (Figure 1B). The single case in which a mutation (3 nucleotide insertion) persisted throughout chronic infection (DK36 clone type) was also the only one that did not cause a frameshift, and thus likely had a lower impact on protein functionality. This suggests that while OprD loss may provide a selective advantage during infection, the trait appears to be disadvantageous for long-term persistence in chronic infection.

### Mutations in oprD increase early attachment and translocation across airway epithelia

The presence of OprD mutations independently of carbapenem treatment suggested that OprD loss may confer infection-related advantages beyond antibiotic resistance. To directly test this hypothesis, we engineered two PAO1 derivative *oprD* mutant strains harboring previously identified mutations ^6^, to avoid the confounding effects of additional mutations present in clinical genomes. One mutant carried a 13-nucleotide deletion (positions 856–868) originally found in a strain with DK41 clone type (*oprD41*) that was selected right after a carbapenem treatment period. The other reconstructed mutant harbored a two-nucleotide (AC) deletion at positions 243–244 previously detected in a clone type DK09 strain (*oprD0S*), that appeared without carbapenem treatment (Figure 1).

We then compared the infection dynamics of these mutants with the wild-type PAO1 strain in a fully differentiated airway epithelium cultured at the air–liquid interface (ALI) which exhibits barrier function and mucus production as a human airway epithelium *in vivo* ^18,21^. Following apical inoculation, bacterial growth, attachment to the epithelial cells and translocation to the basolateral compartment were monitored from 1.5 to 14 hours post-infection.

Both *oprD* mutants displayed consistently higher initial attachment and a markedly increased capacity to cross the epithelial barrier compared with PAO1. CFU counts of bacteria firmly attached to the epithelium (not removed by apical PBS washing) were significantly higher for the mutants than for PAO1 as early as 1.5 hours post-inoculation (Figure 2A, one-way ANOVA with Tukey’s multiple comparisons test, p<0.05 for *oprD*41 and p<0.001 for *oprD*09). Mutants were also detected in the basolateral compartment by 10 hours post-infection, whereas wild-type PAO1 bacteria were absent at this time point, and their numbers remained higher than PAO1 at 14 hours (Figure 2A, one-way ANOVA with Tukey’s multiple comparisons test, p<0.0001). Confocal microscopy analysis confirmed these findings: at 10 hours, mutant bacteria were observed to be already penetrating the epithelium while PAO1 cells remained largely superficial, and by 14 hours, mutants showed substantially deeper and more abundant epithelial penetration (Figure 2B). To assess whether these infection dynamics translated into detectable disruption of the epithelial barrier, we measured transepithelial electrical resistance (TEER) as a general indicator of tight junction integrity and thus overall epithelial damage. TEER progressively declined over the course of all infections, consistent with the large epithelial barrier disruption. However, no significant differences were detected between strains at the monitored timepoints (Figure 2C, two-way ANOVA with Tukey’s multiple comparisons test, *p*>0.05).

**Figure 2.**
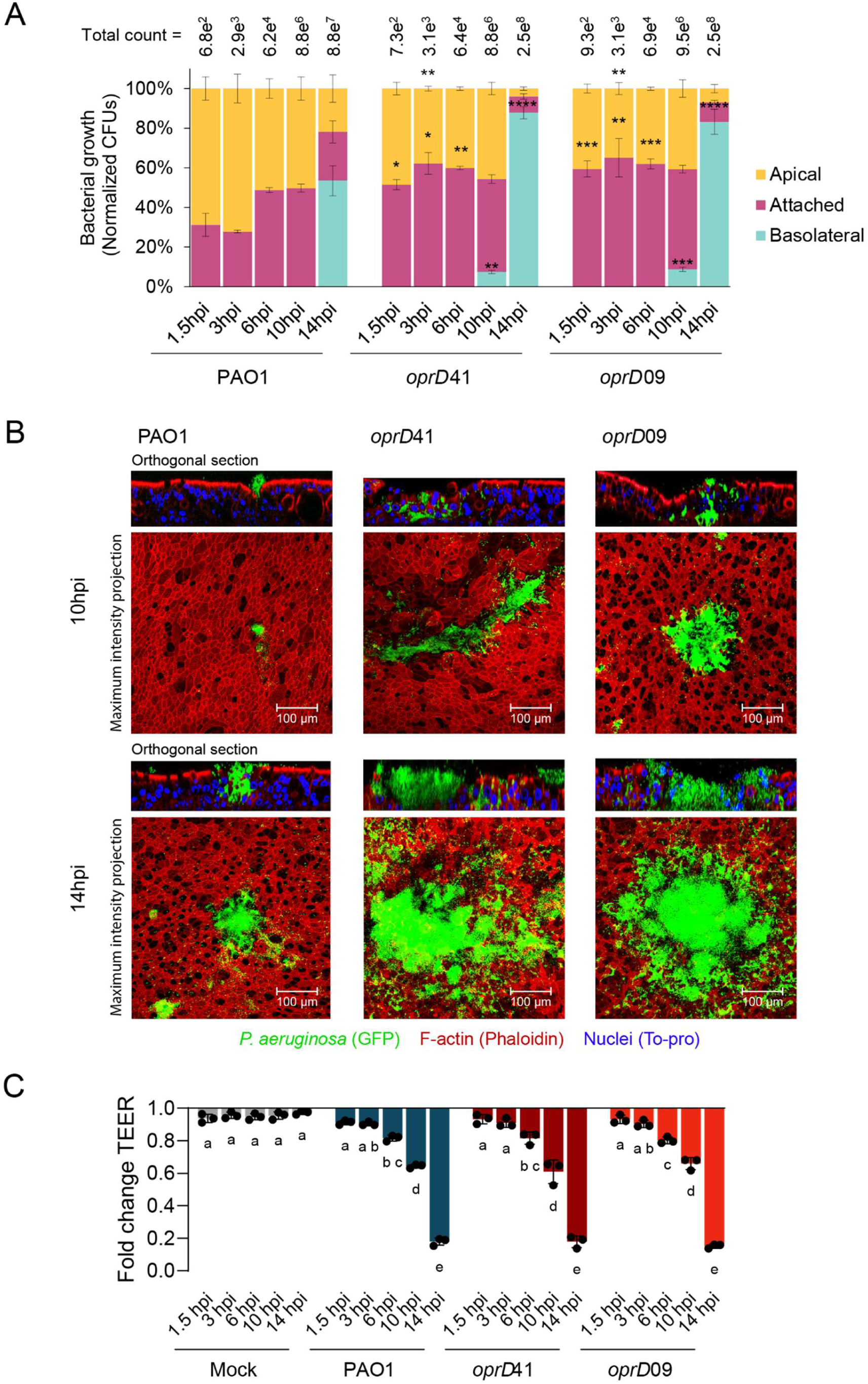
Effects of *oprD* mutations on airway infection dynamics of *P. aeruginosa*. (A) Colony-forming units (CFUs) of PAO1, *oprD*41, and *oprD*09 in the apical compartment (non-firmly attached cells, in yellow), attached to the epithelium (pink), and in the basolateral compartment (blue), measured from 1.5 to 14 hours post-infection in fully differentiated BCi-NS1.1 ALI airway epithelial cultures. Data represent mean ± SEM from three biological replicates, each with three technical replicates. Statistical significance was determined by compartment and timepoint by one-way ANOVA with Tukey’s multiple comparisons test and is indicated as *p ≤ 0.05, **p ≤ 0.01, ***p ≤ 0.001 and ****p ≤ 0.0001. (B) Confocal microscopy images of BCi-NS1.1 ALI cultures at 10- and 14 hours post-infection with PAO1, *oprD*41, and *oprD*09. Bacteria are shown in green (GFP), epithelial F-actin in red (Phalloidin-AF555), and nuclei in blue (TO-PRO-3). Nuclei signal was omitted from maximum intensity projections for image clarity. Scale bar, 100 µm. Images are representative of three independent experiments. (C) Fold change in transepithelial electrical resistance (TEER) of fully differentiated BCi-NS1.1 ALI airway epithelial cultures left uninfected (Mock, in grey) or infected with PAO1 (blue), *oprD*41 (dark red), or *oprD*09 (light red), calculated at the end of the experiment (1.5 to 14 hours) relative to baseline values at the beginning. Data represent mean ± SEM from three biological replicates, each with three technical replicates. Statistical significance was assessed by two-way ANOVA with Tukey’s multiple comparisons test; connecting letters indicate significant differences between groups (*p*<0.05).

These results indicate that loss of OprD enhances *P. aeruginosa* early adhesion to and translocation across the airway epithelium, although this does not affect initial overall tissue disruption.

### Mutations in oprD are associated with changes in membrane organization and adherence to mucin

To investigate the mechanisms underlying the enhanced epithelial crossing of the *oprD* mutants, we performed both phenotypic screenings and transcriptomic analyses on the bacterial monocultures grown in laboratory media Luria–Bertani (LB). In *in vitro* virulence analyses such as pyocyanin production, protease secretion, biofilm formation, and swarming motility, the mutant strains showed no significant differences from the wild-type PAO1 (Supplementary Figure 1A, 1B, 1C and 1D, one-way ANOVA with Tukey’s multiple comparisons test, p>0.05). Likewise, RNA-seq analysis revealed no significant transcriptional changes between strains (Supplementary Figure 2A and 2B), with no genes meeting differential expression thresholds (|log₂ fold change| ≥ 1, adjusted p < 0.05). These results indicate that the absence of OprD does not involve a global transcriptional reorganization.

Because type III secretion (T3SS) has been described as a key determinant of epithelial colonization and penetration ^22^, and its expression is strongly host-induced ^23^, we also quantified the transcript levels of five T3SS genes representing different operons by RT-qPCR from total RNA extracted from infected ALI cultures after 14 hours of infection. Consistent with the RNA-seq data obtained in LB, no significant differences were detected between PAO1 and the *oprD* mutants (Supplementary Figure 2C), supporting that the enhanced epithelial phenotype is not due to altered T3SS expression.

We next assessed bacterial fitness in LB and synthetic cystic fibrosis sputum medium (SCFM) which mimics the nutritional composition of the CF airways ^24^. Growth of the mutants was indistinguishable from PAO1 (Supplementary Figure 1E, one-way ANOVA with Tukey’s multiple comparisons test, p>0.05), ruling out a contribution of general fitness. Antibiotic susceptibility profiling confirmed the expected increase in carbapenem resistance and additionally revealed a modest decrease in colistin susceptibility (Figure 3A, Supplementary Figure 3). Because colistin is an antimicrobial peptide that interacts with the bacterial surface, we hypothesized that OprD loss altered membrane properties.

**Figure 3.**
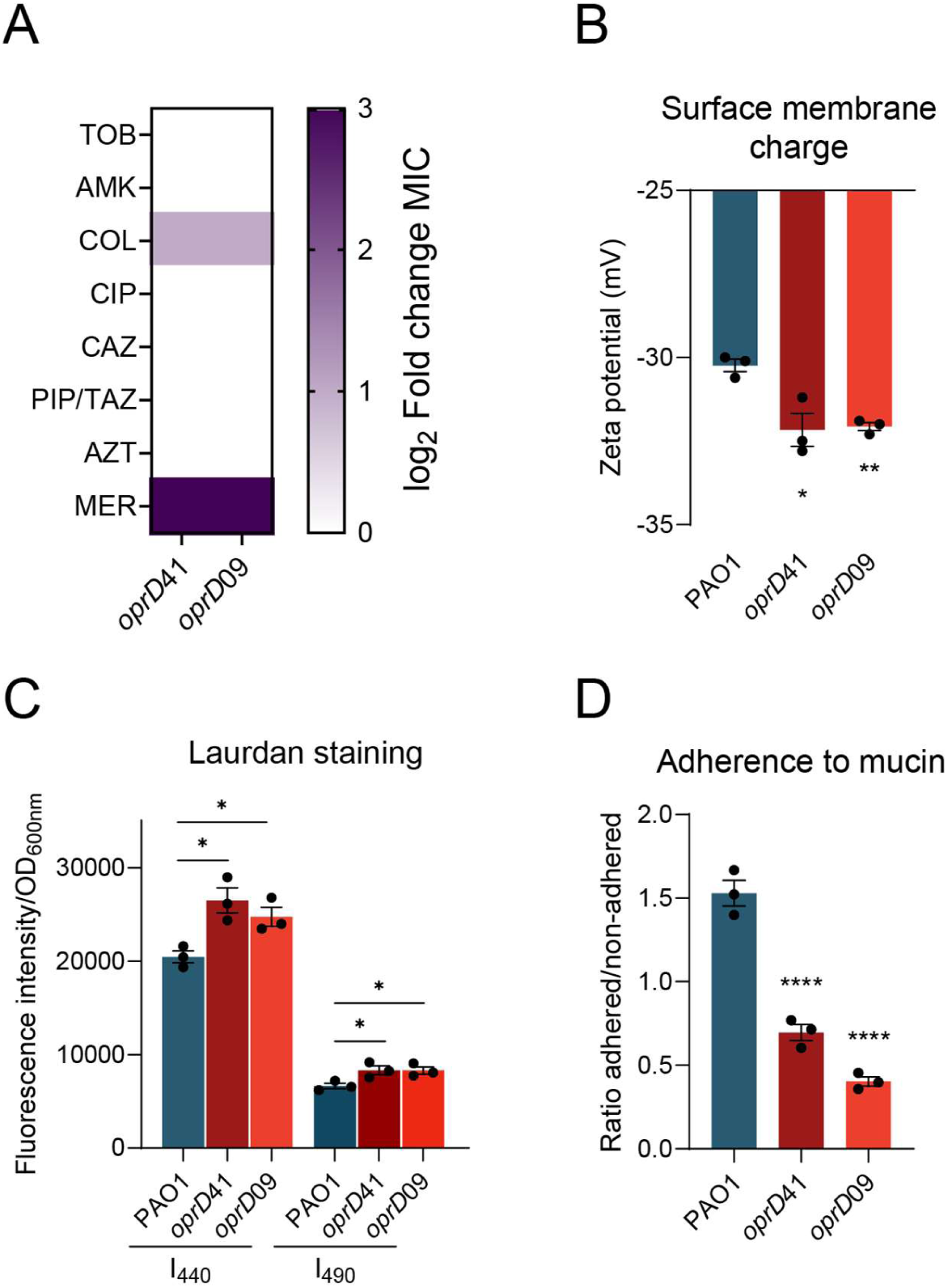
Phenotypic characterization of changes caused by *oprD* mutations in *P. aeruginosa*. (A) Antibiotic susceptibility of *oprD*41 and *oprD*09 relative to the parental PAO1 strain against tobramycin (TOB), amikacin (AMK), colistin (COL), ciprofloxacin (CIP), ceftazidime (CAZ), piperacillin/tazobactam (PIP/TAZ), aztreonam (AZT), and meropenem (MER). Bacterial growth data (OD_600_ values at the end of the assay across antibiotic concentrations) are provided in Supplementary Figure 2. (B) Surface charge and (C) Laurdan fluorescence intensity for PAO1 (blue), *oprD*41 (dark red), and *oprD*09 (light red). Surface charge values are shown as mean ± SEM from three biological replicates with twenty technical replicates each, whereas Laurdan fluorescence total intensity is presented as mean ± SEM from three biological replicates with three technical replicates each. Statistical significance was determined by one-way ANOVA with Tukey’s multiple comparisons test and is indicated as *p ≤ 0.05 and **p ≤ 0.01. (D) Mucin adherence of PAO1 (blue), *oprD*41 (dark red), and *oprD*09 (light red), expressed as the ratio of attached to non-attached bacteria after incubation with mucin. Data represent mean ± SEM from three biological replicates with three technical replicates each. Statistical significance was determined by one-way ANOVA with Tukey’s multiple comparisons test and is indicated as ****p ≤ 0.0001.

Consistent with this hypothesis, we quantified alterations in outer membrane organization. The *oprD* mutants exhibited a significantly more negative net surface charge than PAO1 (Figure 3B, one-way ANOVA with Tukey’s multiple comparisons test, p<0.05 for *oprD*41 and p<0.01 for *oprD*09), as determined by zeta potential measurements, which report on the electrostatic properties of the bacterial surface. To further assess membrane organization, we used the fluorescent probe Laurdan, which is sensitive to lipid packing and allows discrimination between changes in membrane fluidity (generalized polarization, GP) and overall probe accessibility (fluorescence intensity). Laurdan GP values remained unchanged (GP_PAO1_ = 0.51 ± 0.004, GP*_oprD_*_41_ = 0.52 ± 0.002, GP*_oprD_*_09_ = 0.50 ± 0.002, one-way ANOVA with Tukey’s multiple comparisons test, p>0.05), indicating that lipid order and membrane fluidity are preserved. However, total Laurdan fluorescence increased in the mutants (Figure 3C one-way ANOVA with Tukey’s multiple comparisons test, p<0.05), suggesting greater accessibility of outer membrane components to external molecules, consistent with a structural reorganization of the envelope.

We hypothesized that such alterations in surface properties could directly influence how bacteria interact with the host environment. Given that mucus is a highly negatively charged barrier that bacteria must overcome to establish infection ^25,26^, we next examined mucin binding after incubating bacteria with commercially sourced mucin pre-coated onto the wells of a microplate. Non-adherent cells were removed by washing, and the number of bound bacteria was quantified by CFU counting. Notably, *oprD* mutants showed impaired attachment compared to PAO1 (Figure 3D, one-way ANOVA with Tukey’s multiple comparisons test, p<0.0001), which may reduce their entrapment within mucus and thereby facilitate earlier access to the epithelial surface and more rapid penetration through the tissue.

These findings indicate that OprD loss primarily affects membrane properties and mucus interaction, rather than global virulence regulation or bacterial fitness, providing a plausible mechanistic basis for the enhanced epithelial penetration observed.

### Effects of oprD mutations on airway infection are preserved in clinical genomic backgrounds

Finally, to validate the relevance of these findings in patient-derived isolates, we tested whether the infection phenotype was preserved in clinical genomic backgrounds. For that, we analyzed epithelial penetration of pairs of clinical strains with or without *oprD* mutation, isolated from the same sputum sample and phylogenetically related, minimizing the number of confounding genetic differences (Supplementary Table 2). The pairs, previously whole-genome sequenced ^6^, belonged to clone types DK06 (strains 305 and 306) and DK44 (strains 330 and 333). In DK06, strain 306 carried a deletion of a C at position 927 of *oprD* and had emerged after carbapenem therapy (Figure 1), whereas strain 305 retained the wild-type allele. In DK44, strain 330 harbored a two-nucleotide insertion (AC) at position 184, while strain 333 carried an intact *oprD* gene; these isolates were obtained from a patient who had not received prior carbapenem treatment (Figure 1).

Infections were performed in the airway epithelium ALI model, with bacterial growth, attachment and translocation to the basolateral side assessed at 14 and 22 hours post-inoculation. Consistent with the phenotype observed in the PAO1 background, the *oprD* mutant clinical strains (306 and 330) were detected in the basolateral compartment as early as 14 hours, whereas their *oprD* wild-type counterparts (305 and 333) were absent at this time point, and the *oprD* mutant strains were attached to the epithelium in higher numbers (Figure 4A, left panel; t-test with Holm–Šídák multiple comparisons test, p<0.05). By 22 hours, bacteria from all strains were detectable in the basolateral compartment, but *oprD* mutants remained significantly more abundant than their *oprD* wild-type counterparts (Figure 4A, right panel; t-test with Holm–Šídák multiple comparisons test, p<0.05). Confocal microscopy images at 22 hours post-infection visually confirmed these quantitative results, showing deeper epithelial penetration of the *oprD*-mutated strains across distinct colonization patterns (Figure 4B).

**Figure 4.**
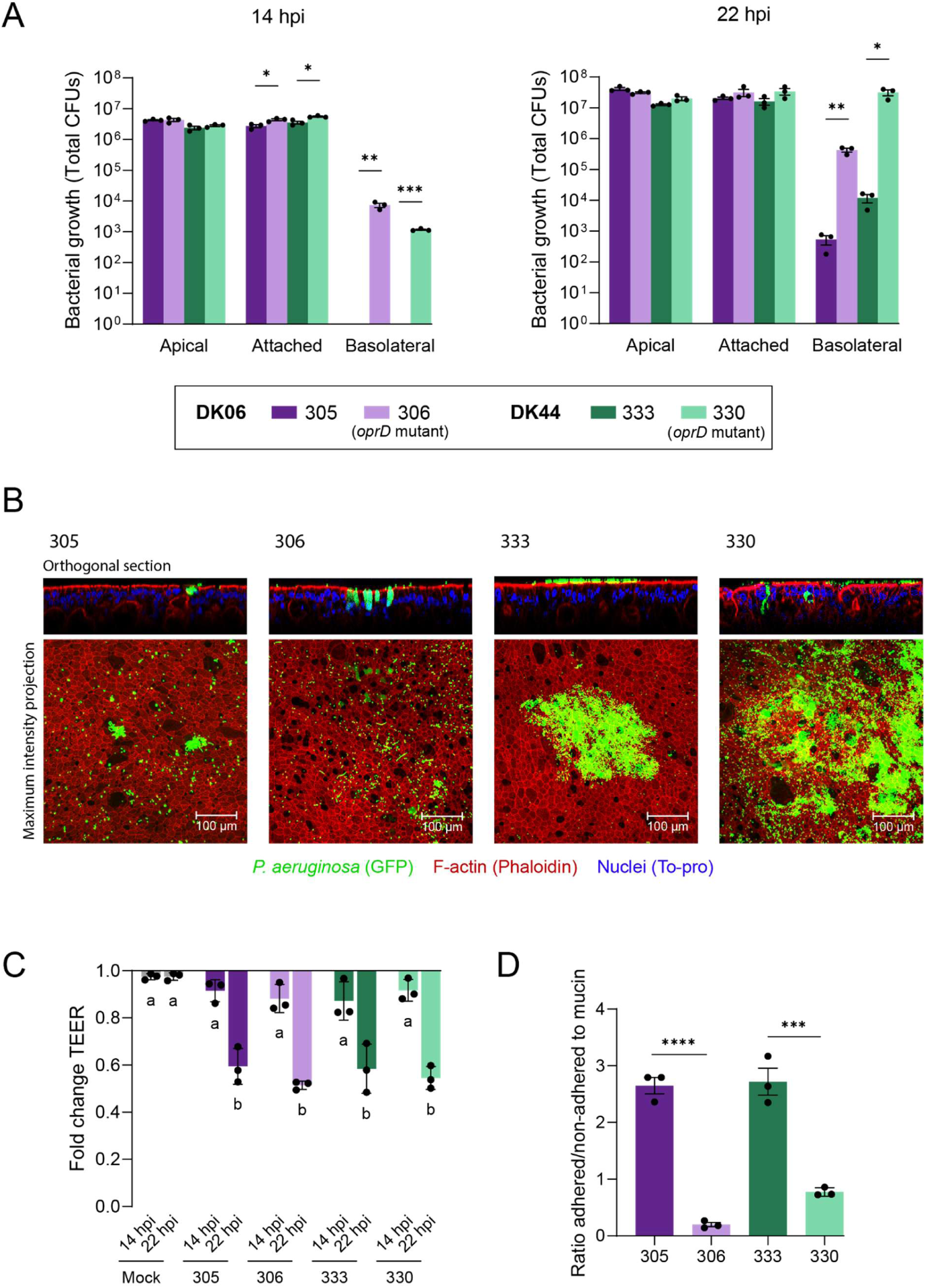
Effects of *oprD* mutations on airway infection dynamics of *P. aeruginosa* clinical strains. (A) Colony-forming units (CFUs) of clinical strains 305 (dark purple), 306 (light purple), 333 (dark green), and 330 (light green) recovered from the apical compartment (non-attached cells), firmly attached to the epithelium, and from the basolateral compartment at 14 (left panel) and 22 hours (right panel) post-infection in fully differentiated BCi-NS1.1 ALI airway epithelial cultures. Data represent mean ± SEM from three biological replicates, each performed in technical triplicates. Statistical significance between strains of the same clone type was assessed by t-test with Holm– Šídák multiple comparisons test and is indicated as *p ≤ 0.05, **p ≤ 0.01 and ***p ≤ 0.001. (B) Confocal microscopy images of BCi-NS1.1 ALI cultures at 22 hours post-infection with clinical strains 305, 306, 333, and 330. Bacteria are shown in green (GFP), epithelial F-actin in red (Phalloidin-AF555), and nuclei in blue (TO-PRO-3). Nuclei signal was omitted from maximum intensity projections for image clarity. Scale bar, 100 µm. Images are representative of three independent experiments. (C) Fold change in transepithelial electrical resistance (TEER) of fully differentiated BCi-NS1.1 ALI airway epithelial cultures left uninfected (Mock, in grey) or infected with clinical strains 305 (dark purple), 306 (light purple), 333 (dark green) or 330 (light green), calculated at the end of the experiment (14 or 22 hours) relative to the values at the beginning. Data represent mean ± SEM from three biological replicates, each with three technical replicates. Statistical significance was assessed by two-way ANOVA with Tukey’s multiple comparisons test; connecting letters indicate significant differences between groups (*p*<0.05). (C) Mucin adherence of clinical strains 305 (dark purple), 306 (light purple), 333 (dark green) and 330 (light green), expressed as the ratio of attached to non-attached bacteria after incubation with mucin. Data represent mean ± SEM from three biological replicates with three technical replicates each. Statistical significance was determined by two-way ANOVA with Tukey’s multiple comparisons test and is indicated as ***p ≤ 0.0005 and ****p ≤ 0.0001.

TEER values, measured as an indicator of epithelial barrier integrity, decreased progressively over time in all strains (Figure 4C; two-way ANOVA with Tukey’s multiple comparisons test, p>0.05 between strains). However, no significant TEER reduction relative to uninfected controls was observed at 14 hours post-infection for any strain (Figure 4C; two-way ANOVA with Tukey’s multiple comparisons test, p>0.05), indicating that these clinical isolates are less damaging to the epithelium than the laboratory strain PAO1 (Figure 2C).

Consistent with the observations in the PAO1 background, the *oprD* mutant clinical strains also showed reduced adhesion to mucin compared with their *oprD* wild-type counterparts (Figure 4D, two-way ANOVA with Tukey’s multiple comparisons test, p ≤ 0.0001), suggesting that loss of OprD similarly decreases mucus entrapment in clinical isolates and may contribute to their enhanced epithelial translocation.

Together, these findings demonstrate that the enhanced epithelial translocation conferred by *oprD* mutations is preserved in clinical isolates with distinct infection dynamics and virulence, underscoring the relevance of this phenotype in patient infection contexts.

## Discussion

Antibiotic resistance is typically only studied in the narrow context of drug susceptibility, yet its broader effects on bacterial physiology and host interactions remain underexplored. Resistance mutations can reconfigure fundamental aspects of bacterial biology, thereby shaping infection outcomes in ways not directly related to antimicrobial therapy. Here, we provide a clear example: loss-of-function mutations in the carbapenem porin OprD in *P. aeruginosa* alter infection dynamics independently of resistance. Specifically, we show that *oprD* mutants exhibit reduced mucin adhesion and enhanced epithelial translocation (Figure 5).

**Figure 5.**
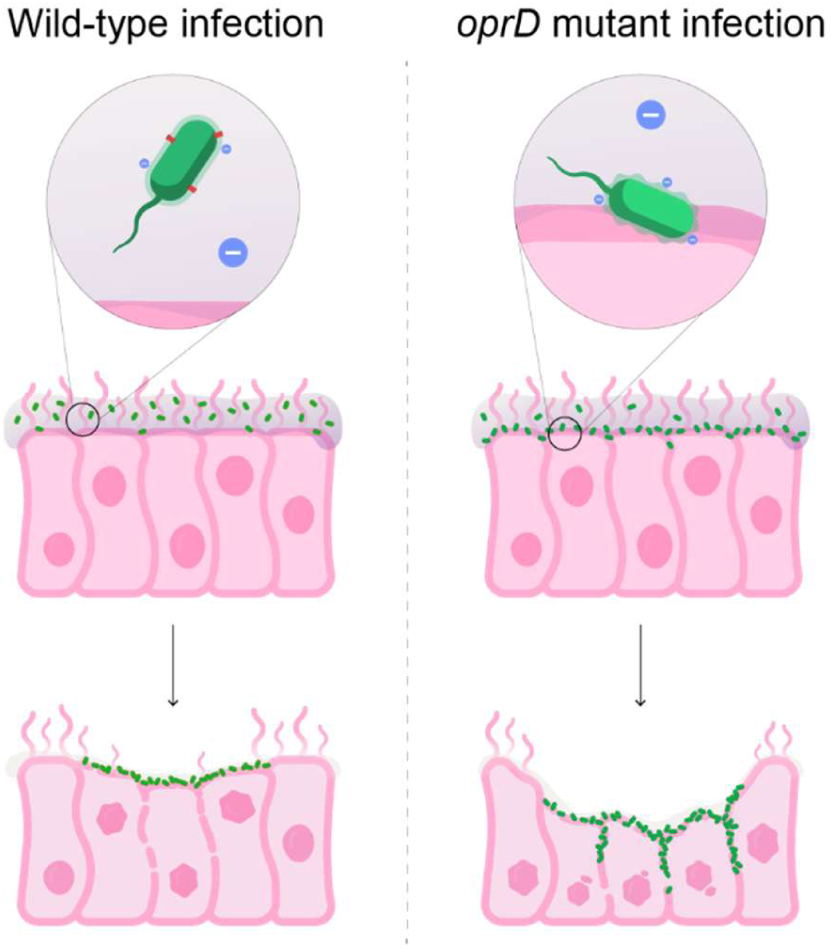
Schematic representation of the effects of *oprD* mutations in *P. aeruginosa* infection. Loss-of-function mutations in *oprD* alter outer membrane organization and increase surface negative charge. These changes likely reduce bacterial binding to mucus, thereby facilitating closer contact with the epithelial surface at early timepoints. This enhanced epithelial interaction promotes more efficient colonization and subsequent translocation across the airway barrier.

This phenotype appears to arise from structural alterations of the outer membrane rather than transcriptional reprogramming. Although direct genetic reversion of only the membrane changes caused by OprD loss is not feasible, and chemical modulation of membrane charge would also affect the host environment, the convergence of our data strongly supports membrane reorganization as the most plausible driver of the observed effects. This interpretation is supported by parallels with *Streptococcus pneumoniae*, whose polysaccharide capsule, particularly when highly negatively charged, reduces mucus-mediated clearance, facilitating transit to and colonization of the epithelial surface ^27^. Notably, *P. aeruginosa* has been shown to breach the airway epithelium via goblet cell invasion ^22^, which requires overcoming entrapment within the mucin secreted by these cells. This suggests that surface modifications reducing mucus entrapment may represent a conserved strategy among respiratory pathogens.

Interestingly, we found that clinical *oprD* mutations can arise both with and without prior carbapenem treatment. Notably, the clinical mutations analyzed in this study, both acquired following carbapenem therapy and in patients without exposure, conferred similar infection-relevant phenotypes. This observation underscores that selective forces acting during infection are not limited to antibiotic exposure. Indeed, previous reports have also described *oprD* mutations emerging in patients without therapy ^28^, further supporting the idea that the selective advantage of *oprD* loss extends beyond drug resistance. One possible explanation is that reduced mucin binding facilitates evasion of mucociliary clearance, providing a selective advantage during airway colonization even in the absence of antibiotic pressure. Importantly, we found that these infection-related phenotypes were maintained across clinical strains with distinct genomic backgrounds, virulence and infection dynamics, indicating that the effect of *oprD* loss is robust and not restricted to a particular strain context. Since the enhanced epithelial penetration observed in *oprD* mutants likely stems from an altered interaction with mucus that promotes earlier contact with epithelial cells, rather than from changes in their colonization strategy, the phenotype can be maintained across strains with different infection dynamics. Nevertheless, we also observed that *oprD* mutations were rarely maintained over time in chronic infections. This transient presence suggests an evolutionary trade-off, as OprD loss facilitates short-term invasion of the airway epithelium but may incur fitness costs, possibly immunological, that limit long-term persistence during chronic colonization. In this sense, *oprD* mutations exemplify the classic evolutionary trade-off in host-pathogen interactions: what benefits acute invasion may ultimately hinder long-term persistence ^29^. Still, it is unclear which other selective pressures may select for *oprD* mutations.

Despite their limited fixation, *oprD* mutations can have clinically relevant consequences. For example, it was reported that, in one CF patient, the acquisition of an *oprD* mutation after carbapenem therapy coincided with a rare increase in virulence during chronic infection ^30^. Although concomitant mutations in the type III secretion system likely contributed substantially to this phenotype, the simultaneous emergence of an *oprD* mutation suggests a possible combined effect. Consistent with this, our data show that the enhanced epithelial penetration associated with *oprD* loss is maintained across clinical isolates with different baseline levels of virulence, as reflected by TEER measurements of the performed infections, indicating that this phenotype is largely independent of strain-specific virulence traits. More broadly, a recent large-scale analysis found that *oprD* mutations were the strongest genomic predictors of septic shock in *P. aeruginosa* infections, outperforming even classical virulence genes ^31^. Together, these findings indicate that *oprD* mutations may shape clinical trajectories in ways not captured by traditional views of resistance.

In addition to their impact on patient outcomes, the enhanced translocation ability of *oprD* mutants may also influence bacterial dissemination. In a murine model of gut infection, *oprD* mutants displayed higher fitness when bacterial recovery from the spleen was used as a readout ^32^, consistent with our mechanistic observations of improved epithelial penetration. Furthermore, in critically ill patients, a well-described gut-to-lung translocation event of *P. aeruginosa* coincided with carbapenem treatment and the emergence of *oprD* mutations ^33^. While these mutations were originally interpreted merely as genetic markers to track strain identity, our results suggest they may actively promote epithelial traversal, facilitating within-host dissemination.

Taken together, our findings reveal that *oprD* loss-of-function mutations promote airway barrier penetration, likely through alterations in outer membrane properties and mucus interaction. However, their lack of stable persistence in chronic infection highlights a trade-off between short-term invasiveness and long-term adaptation. More broadly, our study illustrates that resistance mutations should not be viewed solely through the lens of antibiotic susceptibility. Instead, their interconnection with bacterial physiology can have significant implications for infection dynamics and clinical outcomes beyond the reduction of drug susceptibility. Recognizing these dual roles may ultimately improve how we interpret bacterial genomes in clinical microbiology and refine our predictions of infection progression, with direct implications for patient management.

## Materials and methods

### Bacterial strains growth and antibiotic susceptibility testing

The *P. aeruginosa* laboratory strain PAO1 was employed as the reference background for constructing recombinant strains ^34^. *P. aeruginosa* clinical strains used in this study were previously described and whole-genome sequenced ^6^. The local ethics committee at the Capital Region of Denmark (Region Hovedstaden) approved the use of the stored *P*. *aeruginosa* isolates (registration number H-21078844). Bacterial strains were cultured routinely in LB medium at 37 °C under aerobic conditions.

Bacterial growth was monitored in a BioTek ELx808 Absorbance Microtiter Reader (BioTek Instruments) at 37 °C with shaking at 250 rpm. Optical density (OD) at 600 nm was recorded at 20-minutes intervals over a 24-hours period. Cultures were initiated at an OD_600_ of 0.01, obtained by diluting overnight LB precultures into fresh LB or SCFM. SCFM was prepared following the protocol described previously ^35^, excluding DNA and mucins to reduce viscosity.

The minimum inhibitory concentrations (MICs) of ciprofloxacin, ceftazidime, tobramycin, amikacin, colistin, piperacillin/tazobactam, aztreonam, and meropenem were determined for multiple *P. aeruginosa* strains using the broth microdilution method in Mueller–Hinton (MH) medium, following the guidelines established by the European Committee on Antimicrobial Susceptibility Testing (EUCAST) ^36^.

### Construction of bacterial strains

Genetic modifications were carried out using a CRISPR-Cas9-based recombineering system with a protocol previously described ^37^, that is briefly described below.

Electrocompetent *P. aeruginosa* PAO1 cells were prepared by washing twice with 300 mM sucrose, followed by electroporation with the pSH624-ssr plasmid. Transformants were selected on LB-agar plates supplemented with gentamicin (50 µg/mL). Spacer sequences targeting the *oprD41* mutation (Fw: 5’–GCGCGGGATGCGCACACTTTCACCT– 3’; Rv: 5’–AAACAGGTGAAAGTGTGCGCATCCC–3’) and *oprD0S* mutation (Fw: 5’– GCGCGTATGAATCCGGCTTCACCCA–3’; Rv: 5’–AAACTGGGTGAAGCCGGATTCATAC–3’) were phosphorylated using T4 Polynucleotide Kinase (New England Biolabs) and cloned into Eco31I (FastDigest, Thermo Fisher Scientific)-digested pS548-CsR vector via T4 DNA ligase (Thermo Fisher Scientific), following the manufacturers’ protocols.

To induce ssr expression, an overnight culture of PAO1 pSH624-ssr was incubated with 1 mM IPTG for 3 hours. Subsequently, the constructed pS548-CsR-*oprD41* and pS548- CsR-*oprD0S* plasmids were introduced by electroporation together with the recombineering oligonucleotides containing the desired genetic modification flaked by 50 nucleotides upstream and downstream of its genetic context. Transformants were selected on LB agar containing tetracycline (100 µg/mL), and Cas9 activity was induced by adding 5 mM 3-methylbenzoate (3-mBz) into the LB-agar plate. Successful genome editing was confirmed by PCR screening and Sanger sequencing of the targeted genomic region.

### GFP-tagging of bacterial strains

To enable visualization by confocal microscopy, *P. aeruginosa* strains were genetically modified to express GFP. Tagging was achieved by conjugative transfer of the plasmid pUC18T-mini-Tn7T-Gm-Tp-gfp as previously described ^18,38^. The process involved a four-parental mating, in which equal volumes of cultures of *Escherichia coli* DH5α/pUC18T- mini-Tn7T-Gm-Tp-gfp, *E. coli* SM10(λpir)/pTNS1, *E. coli* HB101/pRK2013, and the target *P. aeruginosa* strain were applied onto a 3-mm cellulose acetate filter placed on non-selective LB agar. Plates were incubated overnight at 37 °C. Transformants were selected on Pseudomonas Isolation Agar (PIA) supplemented with gentamycin (50 µg/mL). Subsequently, the antibiotic resistance cassette was removed by Flp-mediated excision using the plasmid pFLP2. For this step, electrocompetent *P. aeruginosa* cells from the previous conjugation were prepared by two washes with 300 mM sucrose, followed by electroporation with pFLP2. Transformants were selected on LB agar containing carbenicillin (200 µg/mL). Plasmid curing was performed by streaking onto LB agar with 5% sucrose. GFP expression in the final strains was confirmed by detection of green fluorescence.

### Air Liquid Interface infection model preparation

BCi-NS1.1 cells ^21^ were employed to establish ALI-based infection systems as previously described ^17,18^. Cells were first expanded in standard culture flasks under humidified conditions at 37 °C and 5% CO₂, using PneumaCult™-Ex Plus medium (STEMCELL Technologies) according to the manufacturer’s guidelines. For ALI setup, 1×10⁵ cells were seeded onto 1 µm pore polyester membrane inserts (Falcon) pre-coated with type I collagen (Gibco). Once full confluence was achieved, the apical medium was removed, and the basolateral compartment was replenished with PneumaCult™-ALI maintenance medium (STEMCELL Technologies) supplemented with 4 µg/mL heparin, 480 ng/mL hydrocortisone, PneumaCult™-ALI 10× supplement, and PneumaCult™-ALI maintenance supplement. Cultures were maintained for 28 days at 37 °C and 5% CO₂, with media replacement every 3–4 days. Epithelial polarization was monitored by regular measurements of transepithelial electrical resistance (TEER) using an STX2 chopstick electrode (World Precision Instruments). From day 15 onward, accumulated mucus was removed weekly by rinsing the apical surface with PBS.

### Infection procedure and characterization

Overnight cultures of *P. aeruginosa* were diluted in LB medium to an initial OD₆₀₀ of 0.01 and incubated until reaching mid-exponential growth. One milliliter of culture was then pelleted by centrifugation (4500 × g, 5 minutes), washed with PBS, and resuspended in PBS to a final concentration of 1×10⁵ CFU/mL. For infection, 10 μL of this suspension (≈1×10³ CFU) were applied to the apical compartment of fully differentiated BCi-NS1.1 ALI cultures, with 600 μL of supplemented PneumaCult™-ALI maintenance medium in the basolateral chamber. Control wells received the same volume of sterile PBS. Cultures were incubated at 37 °C and 5% CO₂ for the desired infection period, after which 200 μL of PBS were added to the apical chamber.

Apical washes (200 μL) and basolateral media (600 μL) were collected for CFU counting by serial dilution and plating on LB agar. To quantify adherent bacteria, the apical surface was rinsed with 200 μL PBS, epithelial cells were detached using a cell scraper, and resulting suspensions were serially diluted and plated.

For confocal microscopy, infected ALI cultures were rinsed with PBS and fixed with 4% paraformaldehyde applied to both apical and basolateral chambers for 20 minutes at 4 °C. After three PBS washes, samples were permeabilized and blocked with PBS containing 3% BSA, 1% saponin, and 1% Triton X-100 for 1 hour at room temperature. Staining was performed by adding 100 μL of Phalloidin-AF555 (1:400, Invitrogen) and TO-PRO-3 (1:1000, BioLegend) diluted in 3% BSA/1% saponin-PBS to the apical chamber for 2 hours at room temperature. Inserts were then excised, mounted on glass slides with VECTASHIELD® Antifade Mounting Medium (VWR), and imaged using a Leica Stellaris 8 confocal microscope (40× oil objective, NA 1.3). Images were processed with LAS X software v1.4.4.26810 (Leica).

### RNA preparation and RNA-sequencing

Bacteria were inoculated into LB medium at an initial OD₆₀₀ of 0.01 and cultured at 37 °C with shaking at 200 rpm until reaching mid-exponential phase (OD₆₀₀ ≈ 0.6). Three independent biological cultures were assessed for PAO1, *oprD*41, and *oprD*09. Prior to RNA extraction, transcription was blocked by adding RNA Protect Bacteria solution (Qiagen) following the manufacturer’s protocol. Total RNA was then isolated using the RNeasy Mini Kit (Qiagen) according to the supplier’s instructions. RNA quality and integrity were assessed using a Fragment Analyzer 2100 (Agilent Technologies). Samples were run in a NovaSeq Illumina sequencer with 2x150bp configuration yielding 10 million reads per sample.

Reads were trimmed, and low-quality reads and potential contamination from adapters were removed using Trimmomatic (v0.35) tool ^39^. Reads were further processed using SortMeRNA tool (v2.1) ^40^ to remove reads generated from residual rRNA transcripts. Reads were mapped against PAO1 genome (NCBI: NC_002516.2) using BWA-MEM algorithm, and duplicated reads were marked using Picard tools. Reads mapping on each annotated coding sequence were counted using htseq-count v0.7.2 ^41^ imported and processed in RStudio.

Counts were normalized using log2-negative binomial transformation performed using the rld transformation function contained in the R package DESeq2 ^42^ with the option blind set as “true.” Normalized counts were used to evaluate whole transcriptome similarities using hierarchical clustering analysis (HCA), principal component analysis (PCA), and k-mean clustering. DEG analysis between transcriptomes was performed using the R package DESeq2, considering statistically significant genes with a Log2(fold change) ≥ |1| and an adjusted P ≤ 0.01.

### RNA extraction and quantitative RT-PCR from infected cultures

Fully differentiated BCi-NS1.1 ALI cultures were infected with approximately 10³ CFU of each bacterial strain as described above. For each strain, three biological replicates were performed, each consisting of two cell culture inserts pooled to increase bacterial RNA yield. After 14 hours incubation at 37 °C and 5 % CO₂, transcription was stopped by adding 300 µL RNAprotect Bacteria Reagent (Qiagen) per replicate. Total RNA was extracted using the RNeasy Mini Kit (Qiagen) according to the manufacturer’s instructions and treated with TURBO DNase (Invitrogen) to remove residual genomic DNA. cDNA synthesis was performed from 10 µg of RNA with the GoScript Reverse Transcription Mix (Promega). Quantitative PCR was run on a QuantStudio 5 Real-Time PCR System (Applied Biosystems) using 96-well plates and GoTaq qPCR Master Mix (Promega), with 100 ng cDNA per reaction. Cycling conditions were 95 °C for 10 min followed by 40 cycles of 95 °C for 15 s and 60 °C for 1 min. Primers (400 nM) are listed in Supplementary Table 3. Relative expression was calculated by the 2^−ΔΔCT^ method ^43^ using *rplU* as housekeeping reference. Data represents the mean of three biological replicates with three technical replicates each.

### Protease secretion and pyocyanin production

Protease secretion and pyocyanin production were measured as previously performed ^17^.Overnight cultures of *P. aeruginosa* were adjusted to an OD₆₀₀ of 0.01 in 10 mL of LB medium and incubated for 24 hours at 37 °C in 50 mL flasks under constant 250 rpm agitation. Each experiment was performed in three independent biological replicates. One milliliter of each resulting culture was centrifuged (7000 × g, 5 minutes), and the supernatant was sterilized by filtration through a 0.22 μm membrane. Two microliters of the sterile supernatant were spotted onto 1% skim milk agar plates (prepared with 1% agar), and proteolytic activity was evaluated by measuring clearing zones after 24 hours of incubation at 37 °C. The same supernatant was also used for semi-quantitative assessment of pyocyanin production by measuring absorbance at 690 nm in 100 μL aliquots using a BioTek ELx808 Absorbance Microtiter Reader (BioTek Instruments).

### Biofilm production

Biofilm production was assessed as previously described ^44^. *P. aeruginosa* strains were grown in Nunc™ MicroWell™ 96-Well Flat-Bottom Microplates (Thermo Fisher Scientific) containing 150 μL of LB medium per well, with Nunc™ Immuno TSP Lids (Thermo Fisher Scientific) replacing the standard lids. Bacteria were added at an initial OD₆₀₀ of 0.05. Plates were incubated at 37 °C for 20 hours to allow biofilm formation on the pegs. After incubation, the peg lids were rinsed by immersion in a microtiter plate filled with 180 μL PBS to remove non-adherent cells. Subsequently, the lids were transferred to a plate containing 160 μL of 0.01% crystal violet solution for 15 minutes to stain the biofilms. Following staining, the lids underwent three washes in separate PBS-filled microtiter plates (180 μL per well) to eliminate excess dye. Finally, pegs were transferred to a fresh plate with 180 μL of 99% ethanol to solubilize the bound crystal violet. The ethanol solution was then used to quantify biofilm biomass by measuring absorbance at 590 nm in a BioTek ELx808 Absorbance Microtiter Reader (BioTek Instruments).

### Motility assay

Swarming motility was assessed using Petri dishes containing 25 mL of a defined medium composed of 0.5% casamino acids, 0.5% Bacto agar, 0.5% filtered glucose, 3.3 mM K₂HPO₄, and 3 mM MgSO₄, as previously performed ^45^. Four microliters of *P. aeruginosa* cultures adjusted to an OD₆₀₀ of 1.0 were spotted onto the center of each plate. Plates were incubated at 37 °C for 17 hours. Experiments included three technical replicates per strain. Swarming areas were quantified by analyzing photographic images with ImageJ software, measuring the surface area (cm²) covered by bacterial spread.

### Membrane properties determination

Zeta potential measurements of *P. aeruginosa* strains were performed using a Zetasizer NANO-ZS (Malvern Instruments) equipped with a universal dip cell (ZEN1002). Overnight cultures were washed once with ultrapure water to remove residual medium and salts, then adjusted to an OD₆₀₀ of 1. Measurements were obtained from three independent biological replicates, each comprising 30 readings. Electrophoretic mobility values were converted to zeta potentials using the Smoluchowski equation.

Membrane fluidity was assessed via Laurdan staining. Washed bacterial suspensions at OD₆₀₀ = 1 in PBS were incubated with 2.5 µM Laurdan dye for 60 minutes at room temperature in the dark. 150 µL of samples and blanks were aliquoted into Nunc™ 96- Well Optical-Bottom black microplates (Thermo Fisher Scientific) with three technical replicates each. Fluorescence was measured at emission wavelengths of 440 and 490 nm upon excitation at 350 nm in a BioTek ELx808 Absorbance Microtiter Reader (BioTek Instruments). Laurdan GP values were calculated using the formula: GP=(I_440_−I_490_)/(I_440_+I_490_), where I_440_ and I_490_ represent fluorescence intensities at 440 nm and 490 nm, respectively.

### Bacterial mucin adhesion assay

Adhesion assays were conducted as previously performed ^46^ with minor modifications. Wells of Nunc™ MicroWell™ 96-Well Flat-Bottom Microplate (Thermo Fisher Scientific) were coated with 200 µL of porcine mucin (type II, Sigma-Aldrich, M2378) solution (10 µg/mL in PBS), while control wells received 200 µL of PBS. Plates were incubated at 37 °C for 18 hours prior to the assay. Overnight bacterial cultures were washed and diluted in PBS to a final concentration of 5×10^7^ CFU/mL. PBS was aspirated from mucin-coated and control wells, which were then inoculated in triplicate with 100 µL of the bacterial suspension (≈5×10^6^ CFU). Plates were incubated for 30 minutes at 37 °C to allow bacterial adhesion. After incubation, the supernatant was removed, and wells were washed five times with 200 µL PBS to eliminate non-attached cells. The wash solutions were plated in serial dilutions to quantify non-attached bacteria. To detach adherent bacteria along with mucin, 200 µL of 1% (v/v) Triton X-100 in PBS was added to each well and incubated at room temperature for 30 minutes. Serial dilutions of the resulting suspensions were plated to determine CFUs of adhered bacteria. Adhesion efficiency was expressed as the ratio of adhered to non-adhered bacteria. All assays were performed in three biological replicates.

### Statistical analysis

All statistical analyses were performed with GraphPad Prism version 10.3.1 (GraphPad Software). Depending on the experimental design, differences among groups were assessed using ordinary one-way ANOVA or two-way ANOVA followed by Tukey’s post hoc test for multiple comparisons, or by unpaired *t*-tests with Holm–Šídák correction for multiple comparisons. A p-value <0.05 was considered indicative of statistical significance.

### Data availability

All data supporting the findings of this study are available within the paper, its Supplementary Materials, and Supplementary Data files. Raw sequencing data have been deposited in the NCBI Sequence Read Archive (SRA) under BioProject accession number PRJNA1346407. Source data are provided with this paper.

## Acknowledgements

We thank the Infection Microbiology group for their insightful comments and discussion. The Basal Cell Immortalized Non-Smoker 1.1 (BCi-NS1.1) cell line was a gift from Professor Ronald G. Cristal (Weil Cornell Medical College, New York, USA). This research was funded by the “Novo Nordisk Foundation Center for Biosustainability (CfB)” grant number NNF10CC1016517 and by the Challenge Grant NNF19OC0056411 to H.K.J. from The Novo Nordisk Foundation. P.L. is the recipient of an ERS/EU RESPIRE4 Marie Skłodowska-Curie Postdoctoral Research Fellowship (Ref. nr: R4202305-01047; this project has received funding from the European Respiratory Society and the European Union’s H2020 research and innovation programme under the Marie Skłodowska-Curie grant agreement No 847462).

## Author contributions

PL designed the study and performed the experimental work. CAC performed the bioinformatic analysis of transcriptomic data. CAC, RLR, HKJ and SM participated in the design of the study. All authors participated in writing the manuscript and approved the submitted version.

