## Supplementary material for "Carbapenem-resistance *oprD* mutations reshape *Pseudomonas aeruginosa* host-pathogen interactions during infection"

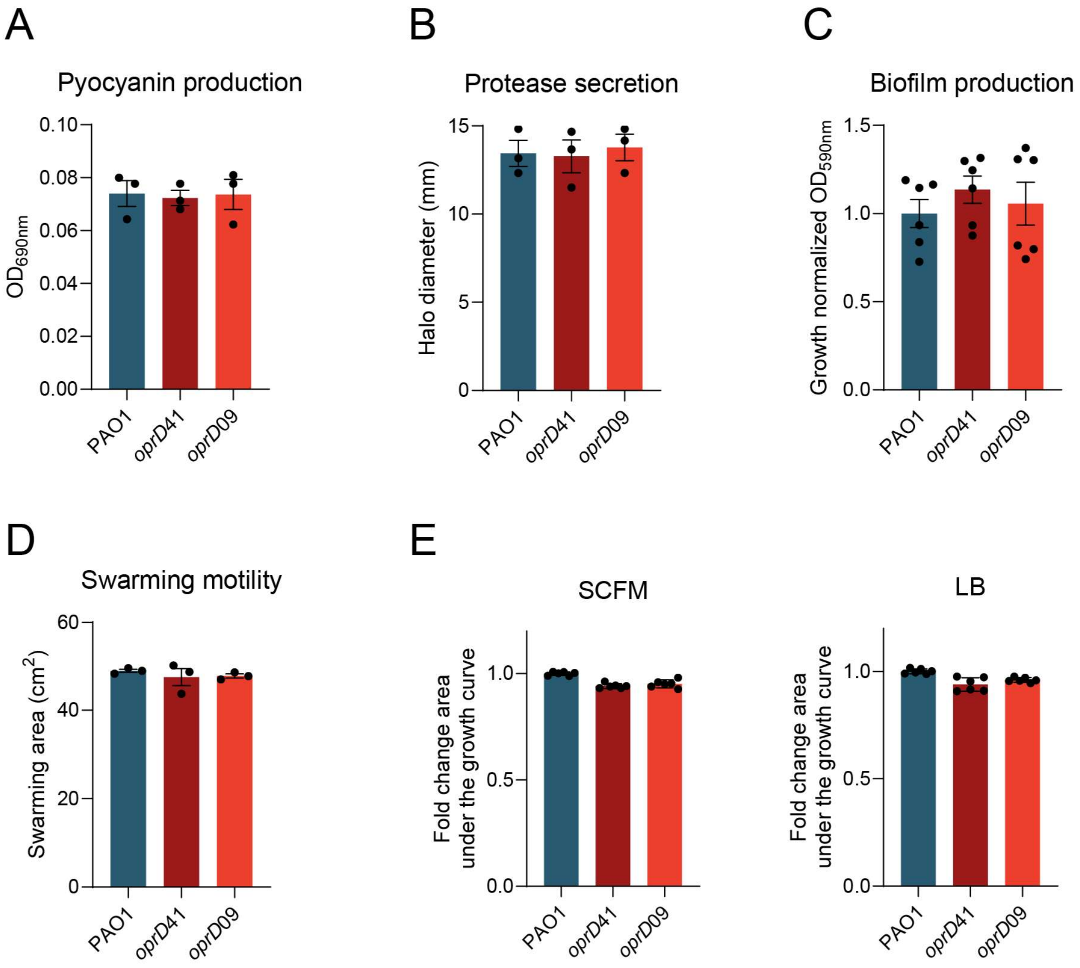


Supplementary Figure 1. Characterization of changes in virulence and growth caused by *oprD* mutations in *P. aeruginosa*. (A) Pyocyanin production, (B) protease secretion, (C) biofilm formation, and (D) swarming motility of PAO1 (blue), *oprD*41 (dark red), and *oprD*09 (light red). Data represent mean ± SEM from three biological replicates with three technical replicates each, except for biofilm formation (six biological replicates). (E) Growth of PAO1 (blue), *oprD*41 (dark red), and *oprD*09 (light red) in LB and SCFM, shown as area under the growth curve. Data represent mean ± SEM from six biological replicates with three technical replicates each.

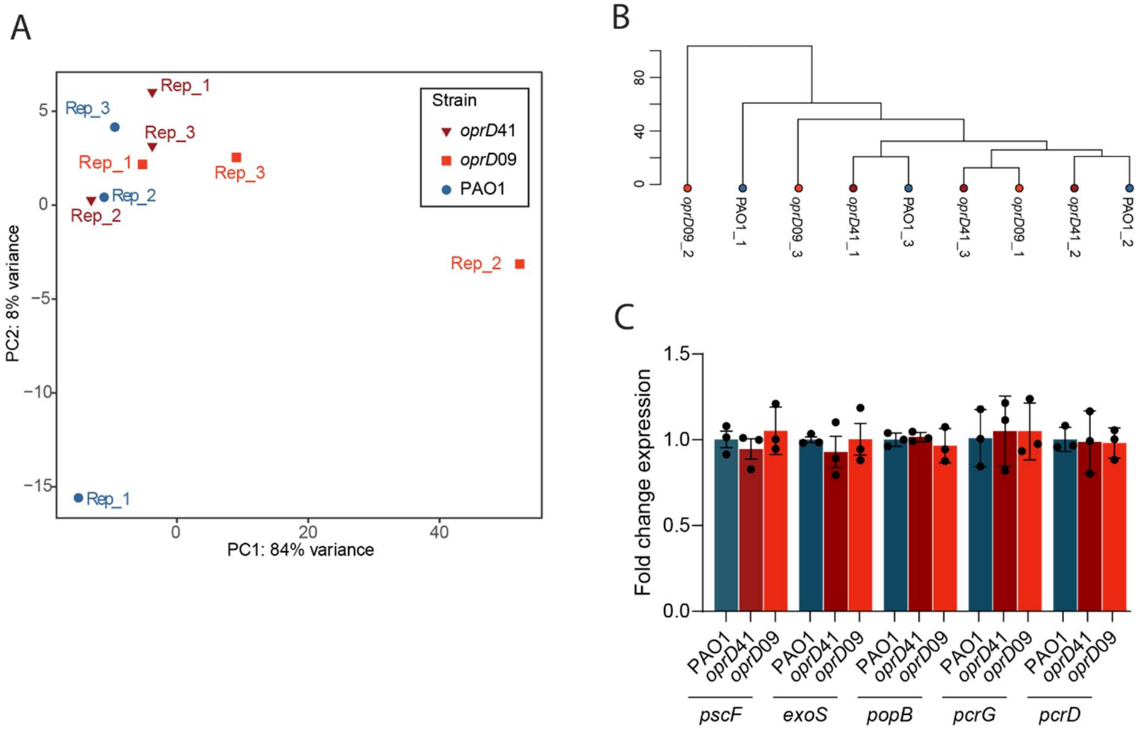


Supplementary Figure 2. Effect of *oprD* mutations on *P. aeruginosa* transcriptome. (A) Principal component analysis (PCA) of variance-stabilized (vsd) count data from 5,678 genes (Source data file). The analysis was performed using the top 500 most variable genes across all samples. Each point represents a sample, colored by strain PAO1 (blue), *oprD*41 (dark red), and *oprD*09 (light red), and the axes indicate the percentage of total variance explained by the first two principal components. (B) Hierarchical clustering dendrogram of the same samples based on Euclidean distances computed from vsd values of all 5,678 genes. The dendrogram illustrates overall similarity among samples, with shorter branch lengths indicating higher transcriptomic similarity. Each sample was run in triplicate and labeled as Rep_1, Rep_2, and Rep_3. DESeq2 differential expression analysis was performed on vsd count data using an absolute $\log_2$ fold change threshold of 1 (lfcThreshold = $\log_2(2)$) and an adjusted *p*-value cutoff of 0.05. No genes were found to be significantly differentially expressed between any sample comparisons under these criteria. (C) Relative expression of *pscF*, *exoS*, *popB*, *pcrG* and *pcrD* in PAO1 (blue), *oprD*41 (dark red), and *oprD*09 (light red) after 14 hours of infection in fully differentiated BCi-NS1.1 ALI airway epithelial cultures, measured by RT-qPCR. Transcript levels were normalized to the housekeeping gene *rplU* to correct for minor differences in bacterial counts and are shown as fold change relative to PAO1, calculated by the $2^{-\Delta\Delta CT}$ method.

Statistical significance versus PAO1 wild type was assessed using one-way ANOVA with Tukey’s multiple comparisons test.

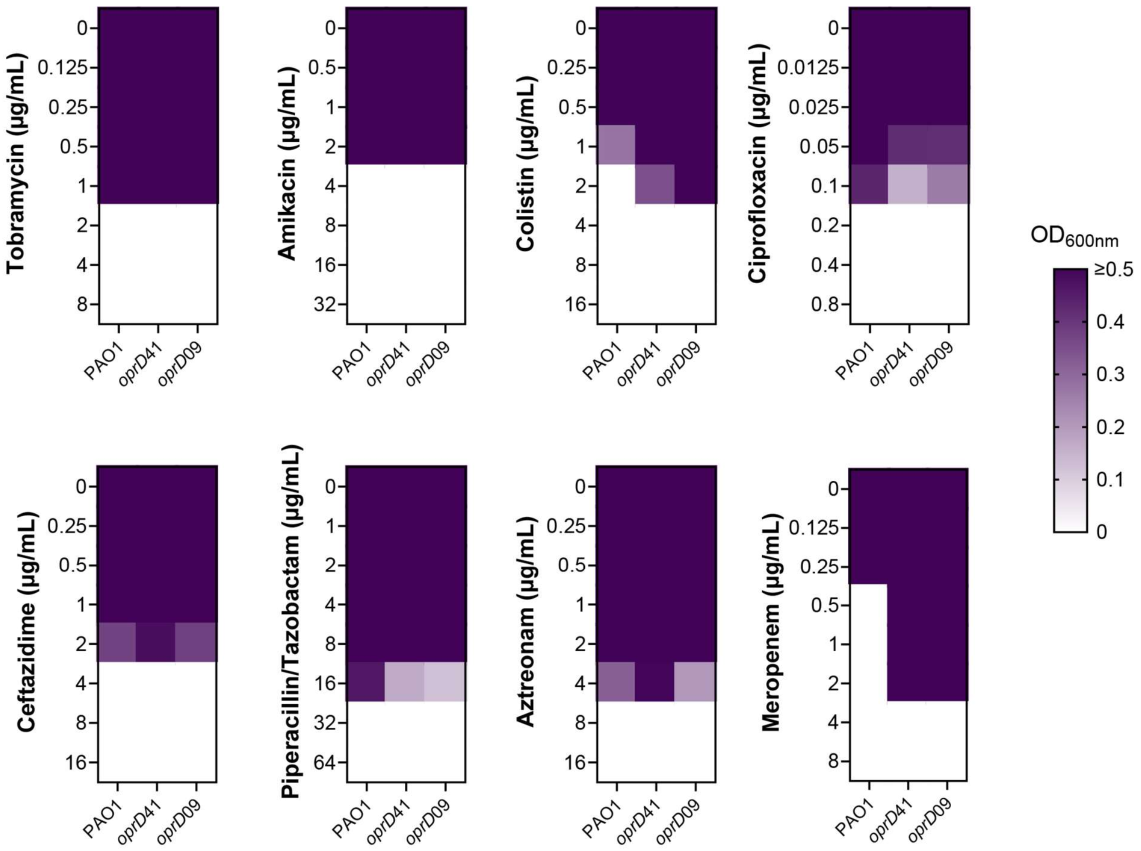


Supplementary Figure 3. Raw antibiotic susceptibility data of wild-type PAO1 and *oprD* mutants. $OD_{600}$ values of PAO1, *oprD*41, and *oprD*09 strains after 24 hours of growth in the presence of increasing concentrations of tobramycin, amikacin, colistin, ciprofloxacin, ceftazidime, piperacillin/tazobactam, aztreonam, and meropenem.

Supplementary Table 1. *oprD*-mutated clinical strains studied in this work and their Minimal Inhibitory Concentration (MIC) for meropenem.

| **Clinical strain ID** | **Clone type** | ***oprD* mutation** | **Meropenem MIC (µg/mL)** |
|---|---|---|---|
| 306 | DK06 | 927 delta(C) | 16 |
| 307 | DK06 | 243 delta(AC) | 24 |
| 82 | DK09 | 243 delta(AC) | 3 |
| 105 | DK12 | 944 delta(GCACCTTCACG) | 64 |
| 107 | DK12 | 944 delta(GCACCTTCACG) | 64 |
| 109 | DK12 | 944 delta(GCACCTTCACG) | 64 |
| 259 | DK35 | 184 ins(AC) | 0.5 |
| 330 | DK44 | 184 ins(AC) | 0.5 |
| 335 | DK44 | 184 ins(AC) | 0.5 |
| 378 | DK41 | 856 delta(GTCTGCGGGAACA) | 12 |
| 399 | DK36 | 184 ins(CTG) | 16 |
| 400 | DK36 | 184 ins(CTG) | 8 |
| 401 | DK36 | 184 ins(CTG) | 512 |
| 403 | DK36 | 184 ins(CTG) | 512 |
| 404 | DK36 | 184 ins(CTG) | 512 |
| 407 | DK36 | 184 ins(CTG) | 512 |
| 402 | DK36 | 184 ins(CTG) | 16 |
| 408 | DK36 | 184 ins(CTG) | 4 |
| 409 | DK36 | 184 ins(CTG) | 16 |
| 397 | DK36 | 184 ins(CTG) | 2 |
| 223 | DK27 | 264 delta(AGTCAGGCCGGGAACGCCATAGG) | 1.5 |
| 15 | DK06 | 104 delta(AC) | 12 |
| 449 | DK12 | 944 delta(GCACCTTCACG) | 1 |

Supplementary Table 2. Genomic differences between paired clinical isolates with and without *oprD* mutations used in this study.

| **DK44 strains** | | | | | |
|---|---|---|---|---|---|
| **Gene locus** | **Gene name** | **Description** | **Mutation** | **Strain 330*** | **Strain 333*** |
| Intergenic PA0819//PA0820 | -//- | 22 downstream hypothetical protein//277 upstream hypothetical protein | 896139-896140 ins(CCCGTTCTCGT) | 1 | 0 |
| PA0958 | *oprD* | Basic amino acid and imipenem outer membrane porin | 1044164-1044165 ins(AC) | 1 | 0 |
| Intergenic PA1334//PA1335 | -//- | 141 upstream probable oxidoreductase//167 downstream probable two-component response regulator | 1447059-1447060 ins(CATTCCCCACA) | 1 | 0 |
| PA2020 | *mexZ* | Efflux pump transcriptional regulators | 2212902-2212921 delta(AACGCCAGGGTGCCGGCGCT) | 1 | 0 |
| Intergenic PA2729//PA2730 | -//- | 622 downstream hypothetical protein//547 downstream hypothetical protein | 3088113-3088123 delta(CCGCATCTCGC) | 1 | 0 |
| PA2904 | *cobI* | Precorrin-2 methyltransferase | 3258918-3258919 ins(GG) | 1 | 0 |
| PA4043 | *ispA* | Geranyltranstransferase | 4526145-4526157 delta(GCCGAGCCCGCCT) | 0 | 1 |

| | | | | | |
|---|---|---|---|---|---|
| Intergenic PA4118//PA4119 | -//*aph* | 17 downstream hypothetical protein//107 downstream aminoglycoside 3'-phosphotransferase type IIb | 4607471-4607472 ins(GCCG) | 0 | 1 |
| Intergenic PA5409//PA5410 | -//*gbcA* | 116 upstream hypothetical protein//159 downstream glycine betaine catabolism protein | 6085226-6085227 ins(CA) | 0 | 1 |

| **DK06 strains** | | | | | |
|---|---|---|---|---|---|
| **Gene locus** | **Gene name** | **Description** | **Mutation** | **Strain 305*** | **Strain 306*** |
| PA0041 | - | Probable hemagglutinin | 53091-53092 ins(GAAA) | 1 | 0 |
| Intergenic PA0366//PA0367 | -//- | 191 upstream probable aldehyde dehydrogenase//99 upstream probable transcriptional regulator | 411125-411125 delta(A) | 1 | 0 |
| PA0720 | - | Helix destabilizing protein of bacteriophage Pf1 | 790477-790478 ins(A) | 1 | 0 |
| PA0888 | *aotJ* | Arginine/ornithine binding protein | 972523-972524 ins(CCCC) | 0 | 1 |
| PA0958 | *oprD* | Basic amino acid and imipenem outer membrane porin | 1044910-1044910 delta(C) | 0 | 1 |
| PA0977 | - | Hypothetical protein | 1060669 G→C | 1 | 0 |

| | | | | | |
|---|---|---|---|---|---|
| Intergenic PA1191//PA1192 | -//- | 45 upstream hypothetical protein//58 upstream conserved hypothetical protein | 1293209-1293210 ins(TGC) | 0 | 1 |
| PA1430 | *lasR* | Transcriptional regulator | 1558838 C→T | 0 | 1 |
| PA2065 | *pcoA* | Copper resistance protein A precursor | 2263646-2263660 delta(GTCCATGCCGTTCAT) | 1 | 0 |
| PA2306 | *ambA* | Aminoacid transporter | 2545339-2545340 ins(TTT) | 1 | 0 |
| PA2443 | *sdaA* | L-serine dehydratase | 2742000-2742001 ins(TT) | 1 | 0 |
| PA2490 | - | Conserved hypothetical protein | 2806291-2806292 ins(ACG) | 1 | 0 |
| Intergenic PA2729//PA2730 | -//- | 140 downstream hypothetical protein//1029 downstream hypothetical protein | 3087631-3087631 delta(T) | 1 | 0 |
| PA3735 | *thrC* | Threonine synthase | 4186790-4186791ins(TTTT) | 1 | 0 |
| PA3789 | - | Hypothetical protein | 4247136 G→A | 0 | 1 |
| PA4000 | - | Hypothetical protein | 4480458-4480458 delta(T) | 1 | 0 |
| PA4092 | *hpaC* | 4-hydroxyphenylacetate 3-monooxygenase small chain | 4575822-4575823 ins(GCGAGC) | 1 | 0 |
| PA4699 | - | Hypothetical protein | 5277279-5277280 ins(C) | 1 | 0 |
| PA5199 | *amgS* | Phosphorilation signal transduction system | 5852455 A→T | 0 | 1 |

| | | | | | |
|---|---|---|---|---|---|
| PA5309 | *pauB4* | FAD-dependent oxidoreductase | 5978719-5978720 ins(CCCC) | 0 | 1 |

*The presence of the respective genetic difference is indicated with a 1 and its absence with a 0.

Supplementary Table 3. RT-qPCR oligonucleotide primers used in this study.

| **Name** | **Sequence (5’→3’)** | **Description** |
|---|---|---|
| exoS_RTPCR_Fw | AGAGAGCGAGGTCAGCAGAG | To quantify *exoS* expression by RT-qPCR |
| exoS_RTPCR_Rv | ATGCCGGTGTAGAGACCAAG | |
| pscF_RTPCR_Fw | CGCACATATTCAACCCCAAC | To quantify *pscF* expression by RT-qPCR |
| pscF_RTPCR_Rv | ATCTTCTGCAGGATGCCTTG | |
| popB_RTPCR_Fw | CTTTGGTTGGATCAGTGCAA | To quantify *popB* expression by RT-qPCR |
| popB_RTPCR_Rv | CCGAGCTTTTCCATCACTTC | |
| pcrG_RTPCR_Fw | CGAATACACCGAAGACACCC | To quantify *pcrG* expression by RT-qPCR |
| pcrG_RTPCR_Rv | CTTGCCACATTTCCGCCAG | |
| pcrD_RTPCR_Fw | GGTGCTGATCGTTTCCATGG | To quantify *pcrD* expression by RT-qPCR |
| pcrD_RTPCR_Rv | TGGATGTTGATCTGCTGGGT | |
| rplU_RTPCR _Fw | CGCAGTGATTGTTACCGGTG | To quantify *rplU* expression by RT-qPCR |
| rplU_RTPCR _Rv | AGGCCTGAATGCCGGTGATC | |
